# Viromes vs. mixed community metagenomes: choice of method dictates interpretation of viral community ecology

**DOI:** 10.1101/2023.10.15.562385

**Authors:** James C. Kosmopoulos, Katherine M. Klier, Marguerite V. Langwig, Patricia Q. Tran, Karthik Anantharaman

**Affiliations:** Department of Bacteriology, University of Wisconsin-Madison, Madison, Wisconsin, USA; Microbiology Doctoral Training Program, University of Wisconsin-Madison, Madison, Wisconsin, USA; Freshwater and Marine Sciences Program, University of Wisconsin-Madison, Madison, Wisconsin, USA; Department of Integrative Biology, University of Wisconsin-Madison, Madison, Wisconsin, USA

**Keywords:** Virome, metagenome, viral ecology, differential abundance

## Abstract

**Background:** Viruses, the majority of which are uncultivated, are among the most abundant biological entities on Earth. From altering microbial physiology to driving community dynamics, viruses are fundamental members of microbiomes. While the number of studies leveraging viral metagenomics (viromics) for studying uncultivated viruses is growing, standards for viromics research are lacking. Viromics can utilize computational discovery of viruses from total metagenomes of all community members (hereafter metagenomes) or use physical separation of virus-specific fractions (hereafter viromes). However, differences in the recovery and interpretation of viruses from metagenomes and viromes obtained from the same samples remain understudied.

**Results:** Here, we compare viral communities from paired viromes and metagenomes obtained from 60 diverse samples across human gut, soil, freshwater, and marine ecosystems. Overall, viral communities obtained from viromes were more abundant and species rich than those obtained from metagenomes, although there were some exceptions. Despite this, metagenomes still contained many viral genomes not detected in viromes. We also found notable differences in the predicted lytic state of viruses detected in viromes vs metagenomes at the time of sequencing. Other forms of variation observed include genome presence/absence, genome quality, and encoded protein content between viromes and metagenomes, but the magnitude of these differences varied by environment.

**Conclusions:** Overall, our results show that the choice of method can lead to differing interpretations of viral community ecology. We suggest that the choice of whether to target a metagenome or virome to study viral communities should be dependent on the environmental context and ecological questions being asked. However, our overall recommendation to researchers investigating viral ecology and evolution is to pair both approaches to maximize their respective benefits.

## INTRODUCTION

Viruses exist in all known ecosystems and infect cells from all domains of life. As the most abundant biological entity on Earth [1,2], viruses significantly impact the ecology and evolution of their hosts [3,4], play pivotal roles in microbial community succession [5], contribute to community-wide metabolic processes [6–8], and are a source of novel therapies being used to combat a worldwide antimicrobial resistance crisis [9,10]. Advances in these areas have been enabled by large-scale investigations into entire communities of viruses which have revealed tremendous amounts of previously unknown virus diversity in human [11–13] and environmental [14–18] systems. Since their hosts largely have not been isolated, these investigations have utilized viral metagenomics (viromics) to examine thousands of viral genomes from DNA/RNA sequence data extracted directly from host-associated and environmental samples. While the number of studies using viromics has been growing in the past decade [18–20], the sampling and analytical methods used vary greatly [20,21]. Although there have recently been efforts to establish standards for analyzing viruses from sequence data [19–21], standards in extraction methodologies are still largely lacking.

There are two ways to identify genomic sequences of viral communities. First, one can sequence metagenomes of a mixed microbial community (hereafter metagenomes). Second, virus-like particles (VLPs) can be separated from a sample to enrich for viral community DNA prior to sequencing (hereafter viromes). Both methods involve computational approaches to identify viral sequences after sequencing, but they each have their own benefits and drawbacks. For instance, viromes do not offer the host context that metagenomes can [22,23]. Thus, investigations into virusLhost relationships can benefit from the use of metagenomes. On the other hand, predicting virusLhost relationships from metagenomes alone remains difficult and can often only be achieved for a fraction of viral genomes [22,23]. Furthermore, rare, low-abundance viruses are diverse and have significant impacts on their communities [24–26]. These viruses are often not detected in metagenomes because viruses represent a small fraction of the mixed community [27]. However, they are detectable in viromes because viruses and other forms of protected environmental DNA represent the majority of sequences in these samples [27,28]. It has also been argued that active viruses exist mostly in an intracellular state and therefore metagenomes are more likely to be appropriate to study viral communities [29,30]. However, the high rates of viral lysis and virion production that have been widely observed [31] might suggest that sequences captured in viromes could better reflect the active viral community. Overall, most studies of viral ecology typically use either method depending on their scope and environmental context.

Although most viral ecology studies have typically utilized either viromes or metagenomes, only a few have leveraged both methods. For example, in an agricultural soil ecosystem, the cumulative richness of viruses in viromes was orders of magnitude greater than that of metagenomes [27]. In a seasonally anoxic freshwater lake, viromes were richer in viruses than metagenomes [6] but the magnitude of this difference was much smaller than that of the soil study. Viral community composition in the freshwater lake was also mostly influenced by sample type (viromes or metagenomes) [6], while human gut viral communities were mostly influenced by the individual human host rather than sample method [32]. These studies offer novel insights into the viral and prokaryotic community composition of their respective ecosystems, but they remain to be synthesized together into a broader context of method application.

The few existing studies that leverage paired viromes and metagenomes have largely paid attention to community-level differences in viruses assembled from each approach, but it remains unknown whether or how this influences the interpretation of ecology and evolution, and the abundance of viruses at the genome level. While differences in genome contiguity and assembly quality between viromes and metagenomes have been discussed [33,34], focused comparisons of viral genomes assembled from viromes versus metagenomes are lacking. Similarly, since the gene content of viruses can vary greatly both within and between populations [35–37], existing community-level comparisons of viromes and metagenomes are unable to highlight any gene-level differences between the two methods.

Here, we directly compare paired viromes and metagenomes from multiple samples obtained from four different environments: a freshwater lake, the global oceans, the human gut microbiome, and soil. After using the same, standardized analytical workflow for every sample and across each environment, we compared viral sequence yields, genome presence/absence, viral genome quality, and virus gene differential abundance between viromes and metagenomes. Last, we discuss the unique insights offered by each approach and suggest when to apply viromes, metagenomes, or both methods when studying viral communities in different environmental contexts.

## METHODS

### Data acquisition

In an effort to compare paired viromes and mixed community metagenomes from a variety of environments, we obtained sequence reads from publicly available studies. We searched for short-read collections that met the following criteria: (1) both viromes and metagenomes must have been generated for the same biological samples, (2) neither virome nor metagenome samples underwent whole-genome or multiple-displacement amplification, and (3) metadata were available that allowed virome and metagenome pairs originating from the same biological sample to be identified, or read filenames made it otherwise clear.

Among the datasets that met the criteria, we chose collections of paired viromes and metagenomes to represent four vastly different environments: a freshwater lake, marine water columns from the global oceans, the human gut microbiome, and soil. Raw reads from virome and metagenome libraries sequenced from water column samples of Lake Mendota, Wisconsin, USA [6] were chosen to represent a freshwater environment. Reads from soil samples of an agricultural field in Davis, California, USA [27] were chosen to represent a soil environment. Fecal sample sequence reads of a cohort in Cork, Ireland [11] were chosen to represent human gut samples. Finally, reads from the Tara Oceans database were obtained to represent marine samples [38,39].

Marine, soil, and human gut reads were obtained from NCBI GenBank [40] using SRAtoolkit (<u>hpc.nih.gov/apps/sratoolkit.html</u>) from BioProjects <u>PRJEB1787</u> (marine metagenomes), <u>PRJEB4419</u> (marine viromes), <u>PRJNA545408</u> (soil viromes and metagenomes) and <u>PRJNA646773</u> (human gut viromes and metagenomes). For the Tara Oceans marine samples, we obtained reads for the <0.22 μm fractions of samples for viromes and the 0.22-3.0 μm fractions for metagenomes (Figure 1A), and read libraries were removed if there was no counterpart library available from the same sample station and depth for the other size fraction. Freshwater virome and metagenome reads were obtained directly by the first author of the study, and can also be found at the JGI Genome Portal under Proposal ID <u>506328.</u> For all environments, all read libraries obtained were composed of paired-end Illumina reads. A detailed description of the data sources for this study and relevant information can be found in Supplementary Table 1.

**Figure 1.**
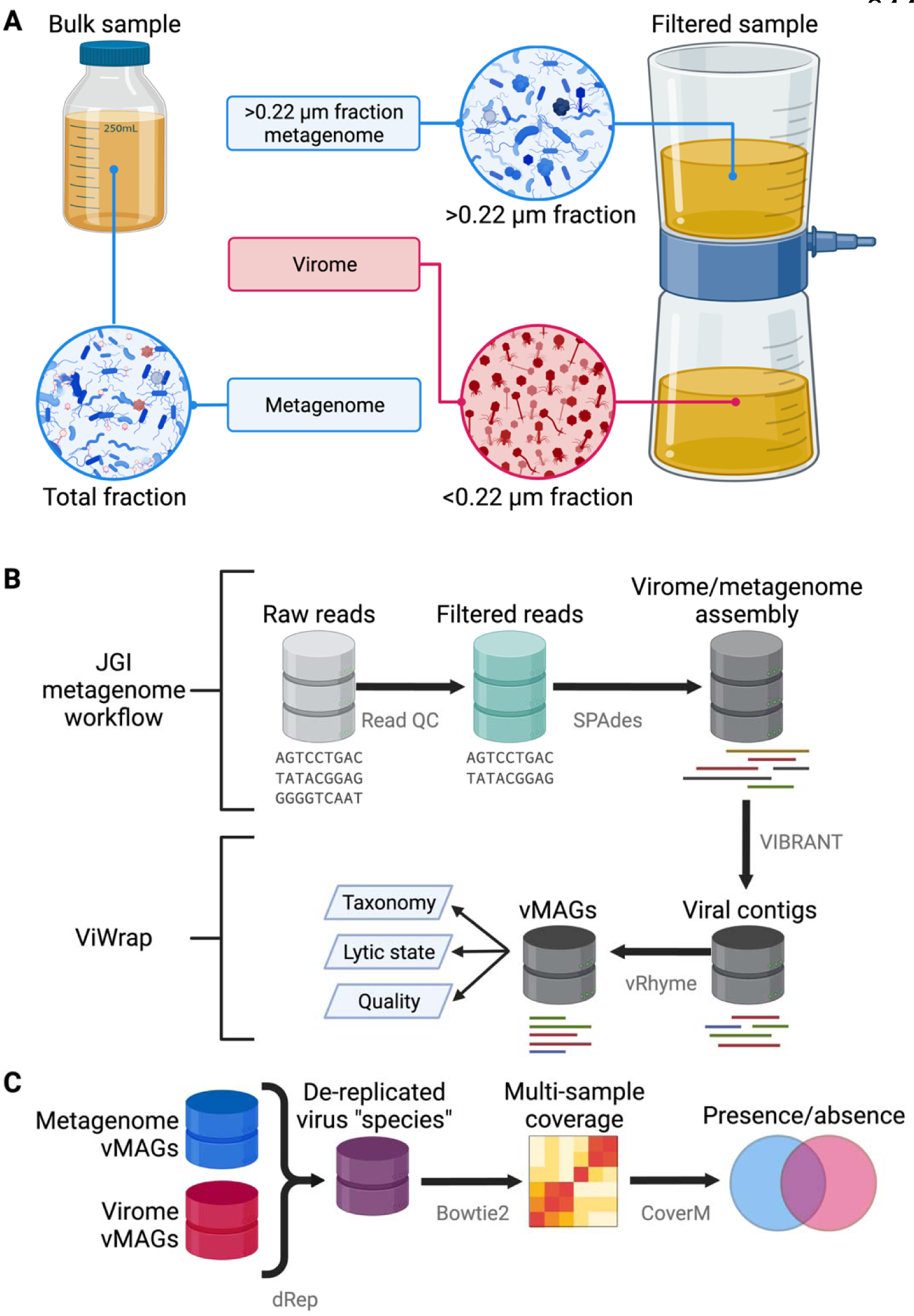
Sampling and analytical approaches used to generate metagenomes, viromes, and vMAGs. (A) Overview of sampling approaches to generate viromes and metagenomes. Viromes were sequenced from a size fraction below 0.22 μm or from a virus-like particle fraction achieved from ultracentrifugation [11,27]. Metagenomes were sequenced using one of two main approaches: DNA from the bulk sample was extracted and sequenced, allowing the recovery of DNA from prokaryotes, viruses, and other microbes. Alternatively, after filtering a sample to isolate virus-like particles in the <0.22 μm fraction, other studies extracted and sequenced DNA from the remaining >0.22 μm fraction that did not pass through the filter [6,38,39]. (B) Overview of metagenome/virome assembly and virus identification methods to obtain viral metagenome-assembled genomes (vMAGs). (C) Overview of methods for the vMAG presence/absence analysis. Figure created with BioRender.com.

### Sequence read quality control and assembly

Freshwater samples were previously sequenced by the Department of Energy Joint Genome Institute (DOE JGI) and thus sequence reads underwent quality control (QC) and were assembled into contigs within the DOE JGI metagenome workflow [41]. To reduce biases that could have been introduced by different QC and assembly methods, read QC and metagenome assembly were performed following the same assembly workflow with the same sequence of software (and versions), commands, and parameters as JGI (Figure 1B). Briefly, raw reads from marine, soil, and human gut samples underwent quality filtering and trimming with BBDuk and BBMap using rqcfilter.sh which were then error-corrected with bbcms. Filtered, error-corrected reads were split into separate mates and singletons using reformat.sh, and the resulting read pairs were imported to metaSPAdes v3.13.0 [42] for assembly. Read lengths and counts at each step of QC were obtained with readlen.sh from the BBTools suite (<u>sourceforge.net/projects/bbmap/</u>) and assembly statistics were obtained for samples from all environments using metaQUAST v5.2.0 [43] which were parsed in R [44] and plotted using ggplot2 [45] to generate Figure 2.

**Figure 2.**
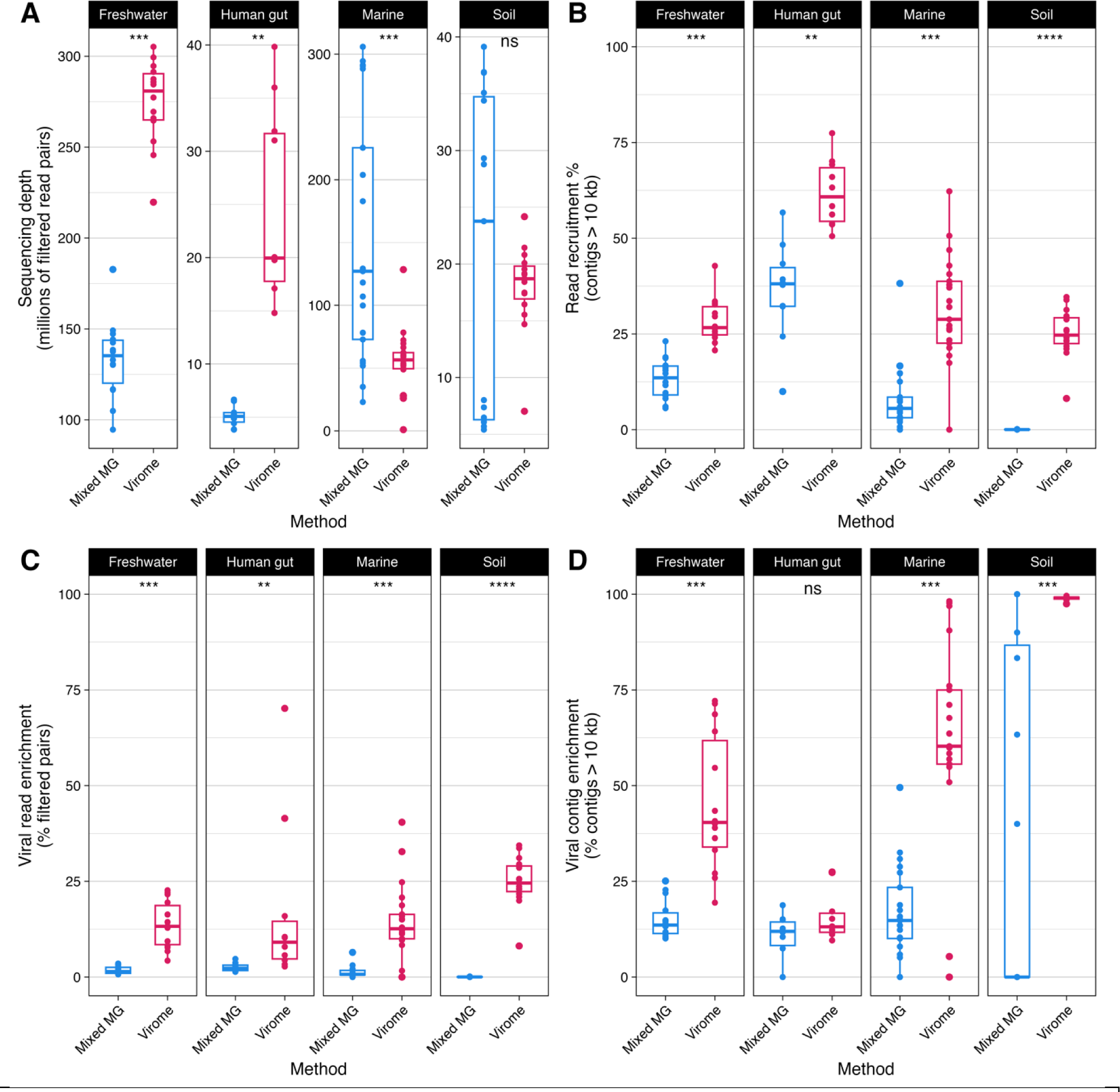
Read recruitment and the enrichment of viral sequences were higher in viromes than metagenomes. Points indicate an individual metagenome/virome assembly. Significance was inferred by Wilcoxon rank sum test: ns *p* >0.05; * *p* ≤0.05; ** *p* ≤0.01; *** *p* ≤ 0.001; **** *p* ≤ 0.0001. (A) While virome samples yielded significantly more read pairs after quality filtering in freshwater and human gut samples, marine metagenomes had greater sequencing depth than viromes, and there was no difference in soil samples. (B) With a minimum alignment identity cutoff of 90%, filtered read pairs from all environments mapped back to assembled contigs >10 kb at a significantly higher rate than metagenomes. (C) In all tested environments, virome assemblies contained more read pairs mapping to viral contigs as a proportion of all quality-filtered read pairs (mapped or unmapped) than metagenome assemblies. (D) All tested environments except human gut samples contained a greater proportion of viral contigs to all assembled contigs >10 kb.

### Virus identification, mapping, binning, quality assessment, and taxonomic assignment with ViWrap

For every sample, ViWrap v1.2.1 [46] was run (Figure 1B) with the assembled sample contigs and filtered reads using the parameter “--identify_method vb” to only use VIBRANT v1.2.1 [47] to identify viral contigs, as well as the options “--input_length_limit 10000” and “--reads_mapping_identity_cutoff 0.90” to adhere to established recommended minimum requirements for virus detection [20]. In accordance with these standards for virus detection, only viral contigs of at least 10 kb were retained for downstream analyses. After using VIBRANT to identify viral contigs, ViWrap mapped reads to the input assembly using Bowtie2 v2.4.5 [48]. Read recruitment to all assembled contigs at least 10 kb was calculated using SAMtools v1.17 [49] using the read mapping files generated by Bowtie2. Read recruitment statistics were then filtered to only include the viral contigs with a length of at least 10 kb identified by VIBRANT. Additionally, ViWrap used the resulting coverage files to bin viral contigs into vMAGs with vRhyme v1.1.0 [50].

In this study, both binned viral contigs and unbinned singletons are together referred to as vMAGs. The quality, completeness, and redundancy of the resulting vMAGs were assessed with CheckV v1.0.1 [51] by ViWrap. ViWrap then grouped vMAGs within samples into genus-level clusters with vConTACT2 v0.11.0 [52] and then into species-level clusters with dRep v3.4.0 [53]. ViWrap assigned taxonomy to vMAGs by aligning proteins with DIAMOND v2.0.15 [54] to NCBI RefSeq viral proteins [55], the VOG HMM database v97 [56], and IMG/VR v4.1 high-quality vOTU representative proteins [57]. Summary statistics on the number of viral contigs, read recruitment, vMAGs, taxonomy, and genome quality gathered by ViWrap for each sample were parsed in R and plotted using ggplot2 to generate Figure 2, Figure S2, Figure S3, and Figure S4.

### Predicting the lytic state of vMAGs

ViWrap provides a prediction of the lytic state for all vMAGs it identifies [46], i.e., whether a vMAG is likely to represent a lytic virus, a lysogenic virus, an integrated prophage flanked with cellular DNA, or not determined. ViWrap makes these determinations based on a combination of annotation results from VIBRANT and binning results from vRhyme. Possible predictions by ViWrap include “lytic scaffold”, “lytic virus”, “lysogenic scaffold”, “lysogenic virus”, and “integrated prophage”. ViWrap handles instances when vRhyme bins multiple integrated prophage sequences or lytic and integrated prophage sequences together by splitting the vMAG back into individual scaffolds to avoid retaining potentially contaminated bins (see <u>github.com/AnantharamanLab/ViWrap</u>). Furthermore, the distinction made by ViWrap between “scaffold” and “virus” depends on the genomic context of the contigs in a vMAG [50] and the estimated completion of a vMAG [51]. Here, we simplified these predictions using a custom python script and did not distinguish between predictions on the “virus” or “scaffold” level and used the results predicted by ViWrap to label vMAGs as “lytic”, “lysogenic”, or “integrated prophage”.

### vMAG presence/absence analysis

Although ViWrap employed dRep to dereplicate vMAGs into species-level clusters at 95% ANI within samples, species representative vMAGs were still redundant between samples after running ViWrap on each. To dereplicate vMAGs across all samples, an additional ANI-based approach was taken. Redundant vMAGs from each sample were gathered and dereplicated using dRep v3.4.3 [53] with a minimum genome length of 10 kb in addition to the options “-pa 0.8 -sa 0.95 -nc 0.85” to set the ANI thresholds for primary and secondary clusters to 80% and 95%, respectively, and to require a minimum covered fraction of 85%, as recommended by established benchmarks for viral community analyses [20]. The parameters “-comW 0 -conW 0 -strW 0 -N50W 0 - sizeW 1 -centW 0” were also used when running dRep so the resulting species representative vMAGs were simply the largest vMAGs in each cluster.

Bowtie2 mapping indices were created from fasta files containing all representative vMAGs from each environment, separately, to be used in competitive alignments. For each environment, filtered reads from every sample were separately mapped to the environment’s mapping index using Bowtie2 v2.5.1 with default parameters to perform an end-to-end alignment and report single best matches at a minimum of 90% identity. The resulting alignment files were sorted and indexed using SAMtools v1.17 [49]. Sorted and indexed files were used with CoverM v0.6.1 (<u>github.com/wwood/CoverM</u>) to obtain covered fraction (genome breadth) statistics at the vMAG level for reads mapping with at least 90% identity. A minimum breadth threshold of 75% was used to establish the detection of a vMAG in each read sample in accordance with previously established recommendations [20]. Lists of unique representative vMAG IDs determined to be present in samples in this way were used to generate Figure 3 and Figure S4 with the R package eulerr (<u>CRAN.R-project.org/package=eulerr</u>) [58,59]. Labels for Figure 3 were manually edited for clarity.

**Figure 3.**
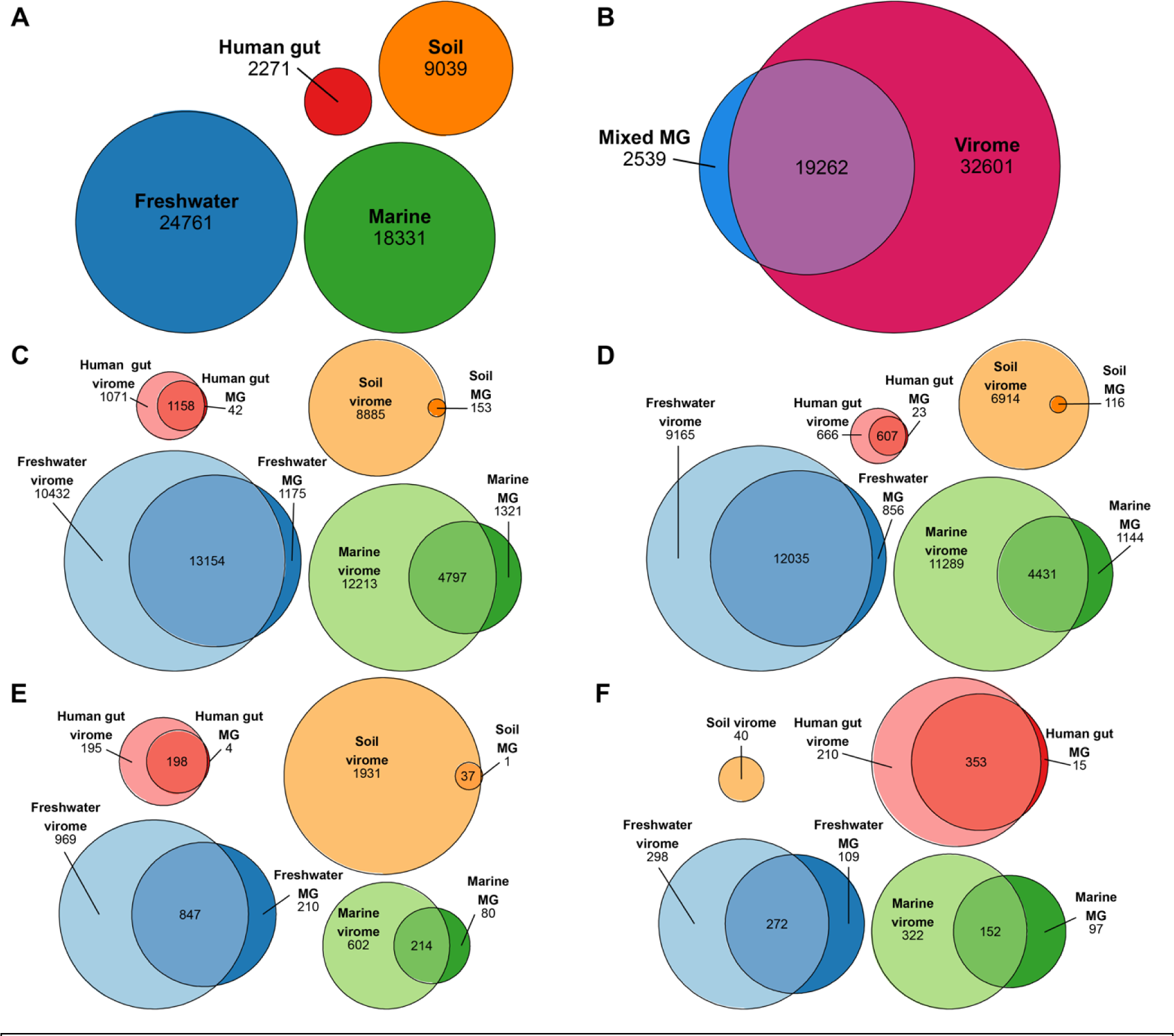
vMAGs assembled from viromes were not detected in most metagenome samples. Euler diagrams generated using eulerr (CRAN.R-project.org/package=eulerr) [58,59] with IDs of unique species-level vMAGs detected in the labeled category; quantities within areas are given beneath labels. An individual vMAG was marked as detected in a virome/metagenome if reads from the virome/metagenome mapped to the contigs in the vMAG with a minimum breadth of 75% across the entire vMAG. (A) Total number of vMAGs in each environment, regardless of method. (B) All vMAGs and environments, separated by method. (C) All vMAGs, separated by environment and method. (D) Predicted lytic vMAGs, separated by environment and method. (E) Predicted lysogenic vMAGs, separated by environment and method. (F) Predicted integrated prophage vMAGs, separated by environment and method.

### Virus genome assembly comparison

To address a preexisting notion that metagenomes typically result in truncated or less-complete viral genome assemblies than viromes [21,27,60], we identified vMAGs shared between viromes and metagenomes. Using our previously generated dRep results, we identified pairs of vMAGs that met the following criteria: (1) one vMAG was assembled from a virome and the other a metagenome, (2) each vMAG in the pair was placed in the same species-level cluster, (3) both vMAGs were assembled from the same sample source, (4) the virome-assembled vMAG was a single contig and predicted by CheckV to be complete, and (5) the metagenome-assembled vMAG was predicted by CheckV to be incomplete.

A single pair was chosen among the resulting candidates based on their respective lengths. Each genome was then subjected to noncompetitive mapping of filtered reads from the virome and metagenome of the same sample source. This resulted in four read mapping files: virome reads mapped to the virome-assembled vMAG, virome reads mapped to the metagenome-assembled vMAG, metagenome reads mapped to the virome-assembled vMAG, and metagenome reads mapped to the metagenome-assembled vMAG. For each file, the read depths d at each genome position were obtained using SAMtools v1.17 [49] with the option “depth”, and then log_10_ normalized by the total number of reads in the sample n in hundreds of millions to obtain a normalized read depth.

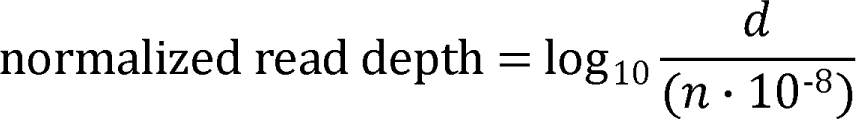

The two vMAGs were aligned using Mauve [61] and BLASTn v2.5.0 from the BLAST+ suite [62] to identify regions in the virome-assembled genome that were missing from the metagenome-assembled genome, as well as gaps and alternate sequences. This revealed the metagenome-assembled vMAG in the pair to be on the opposite strand as the virome-assembled vMAG, so downstream analyses of this vMAG were performed on its reverse-complement. Finally, each vMAG in the chosen pair was reannotated for gene predictions and function using Pharokka v1.4.1 [63] with default settings. The resulting read depths by genome position and unassembled regions were plotted using ggplot2 and arrows representing gene prediction coordinates were added with gggenes v0.5.1 (<u>wilkox.org/gggenes</u>) to generate Figure 4. Highlighted regions and coloring for a selection of genes of interest were added manually to Figure 4.

**Figure 4.**
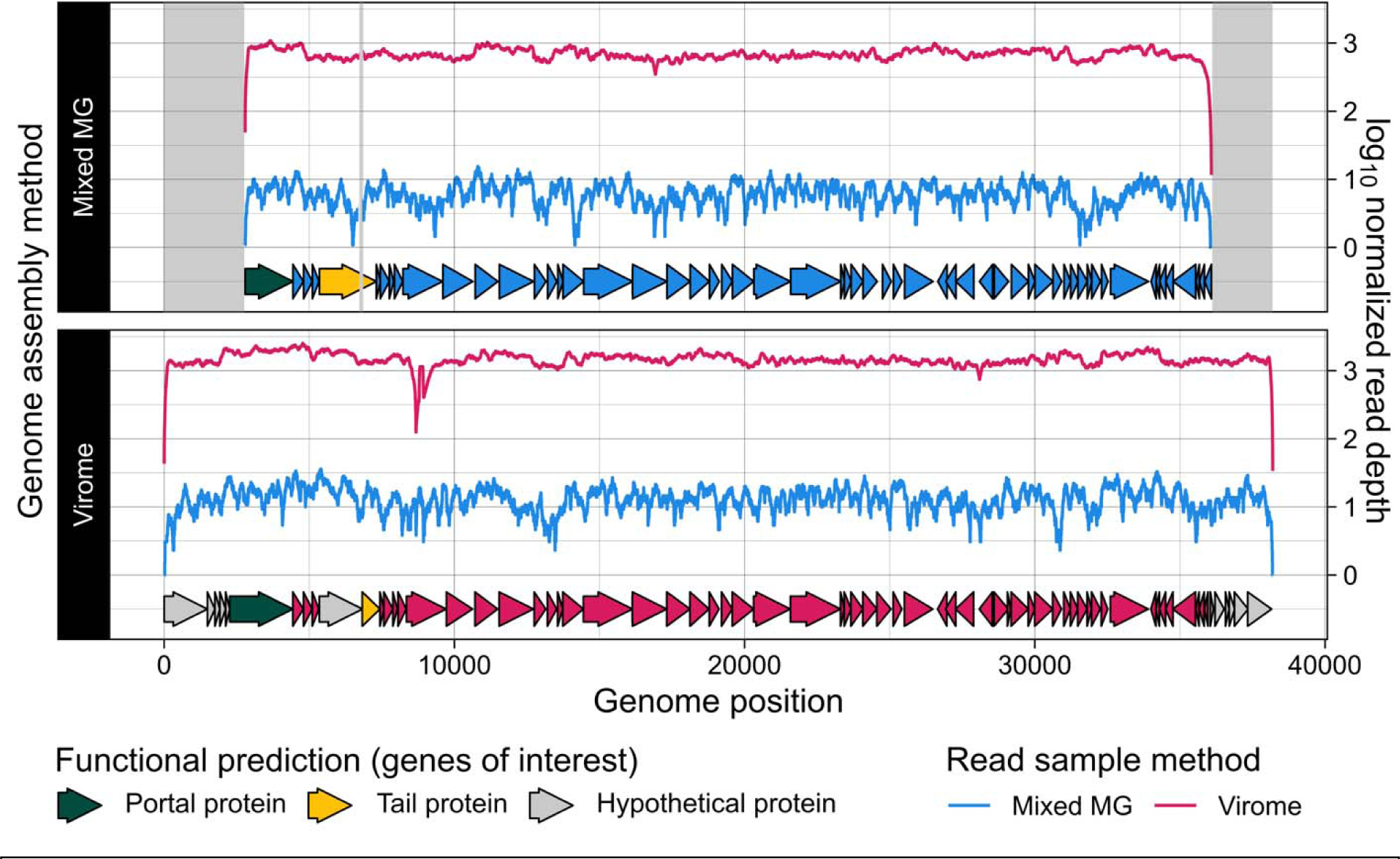
An incomplete metagenome-assembled viral genome was complete in its corresponding virome. A single-contig, complete viral genome identified from a virome assembly was detected but was incompletely assembled in the sample’s corresponding metagenome. Areas highlighted in gray represent regions in the virome-assembled genome that were absent from the metagenome-assembled genome. Reads yielded from the virome and metagenome of the same sample source were each mapped to both versions of the genome assembly. Arrows along the x-axis represent predicted genes that are colored by the extraction method of their genome’s origin, except for a selection of genes of interest that are colored by their functional predictions.

### Differential abundance of viral proteins

We sought to identify protein-coding viral genes that were differentially abundant across virome and metagenome assemblies. For each environment (both viromes and metagenomes), we combined all nucleotide sequences of protein-coding genes predicted by Prodigal [64] that were encoded on viral contigs >10 kb identified by VIBRANT into a database of redundant gene sequences. These databases were then dereplicated, separately by environment, using MMseqs2 v14.7e284 [65]. We used the command “mmseqs easy-search” to estimate pairwise average nucleotide identities (ANI) for all genes in each database, with parameters “--min-seq-id 0.95 -c 0.80 –cov-mode 1” to only retain alignments with minimum ANI of 0.95 and a minimum aligned fraction to the target sequence of 0.80. A clustered graph was generated from the pairwise ANI estimates using mcl with mcxload v14-137 [66] to obtain gene clusters, and the longest gene within each cluster was chosen to be the cluster’s dereplicated representative. Bowtie2 mapping indices were separately generated from the four databases of dereplicated gene representatives of each environment. For each environment, filtered reads from all samples were mapped to the Bowtie2 index of dereplicated genes corresponding to the same environment, using the same parameters and filtering steps as in the vMAG presence/absence analysis above.

Tables of raw mapped read counts for each dereplicated gene representative were obtained for each environment using CoverM. These tables were used to build negative binomial generalized models of gene counts with DESeq2 [67] to infer genes that were differentially abundant across viromes and metagenomes for each environment, separately. The extraction method (virome or metagenome) and sample source were included as factors in the models for each environment, and the DESeq2 workflow employed Wald tests to compare the counts between viromes and metagenomes. For each test, the resulting log_2_ fold changes reported by DESeq2 were shrunken using the function “lfcShrink” with adaptive Student’s *t* prior shrinkage estimators. We used a false-discovery rate adjusted *P*-value cutoff of 0.05 for the Wald test results as well as a minimum shrunken log_2_ fold change of 0.58 (corresponding to a minimum fold change of 1.5) as requirements to determine if a given gene was enriched in either virome or metagenome samples of a given environment. The results were visualized using ggplot2 to generate Figure 5A.

**Figure 5.**
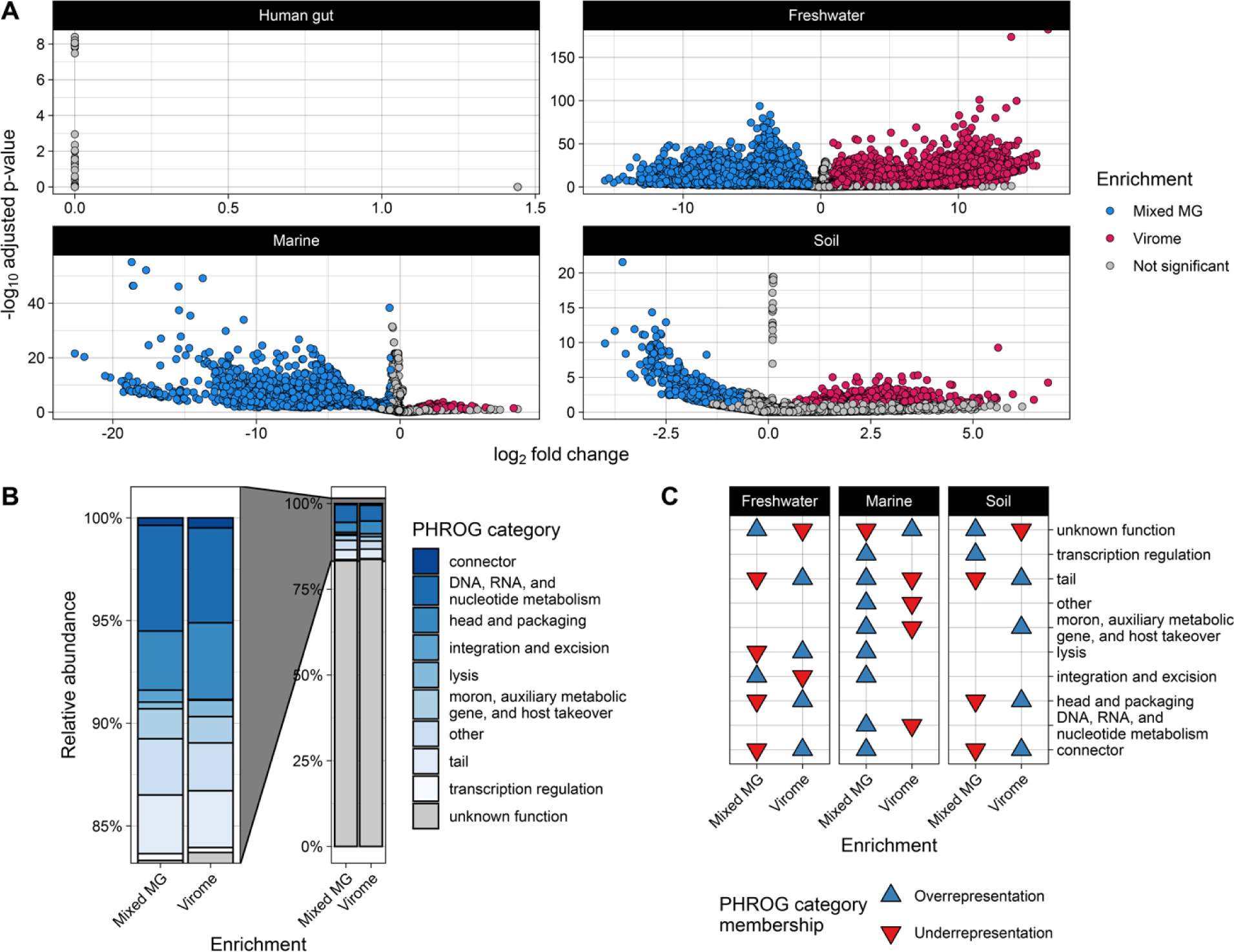
Protein-coding viral genes are differentially abundant across viromes and metagenomes and have predictable functions. (A) Differential abundance of protein-coding viral genes as inferred by DESeq2 [67]. Points indicate unique, dereplicated protein-coding viral genes that were annotated from viral contigs assembled from the environment indicated by the panel labels. Enrichment of a given gene in virome or metagenome samples was determined if the resulting fold change was at least 1.5. (Wald test *P* <0.05, FDR adjusted). No protein-coding viral genes were determined to be significantly enriched in the virome or metagenome human gut assemblies. (B) Relative abundance and (C) over/underrepresentation of PHROG [68] functional categories assigned to differentially abundant genes displayed in (A) (hypergeometric test *P* <0.05, FDR adjusted). Categories without an arrow in a given environment/method were not significantly over or underrepresented in that environment/method.

PHROG [68] functional predictions for all dereplicated gene representatives were obtained by running Pharokka v1.4.1 [60] on each dereplicated gene database. The resulting PHROG annotations and functional categories were mapped back to the DESeq2 significant genes to obtain the presence of PHROG functional categories in each enrichment (virome or metagenome). The relative abundance of PHROG categories among all genes in each enrichment group was calculated and plotted with ggplot2 to generate Figure 5B. To assess the over- or underrepresentation of any PHROG category within either enrichment group, we performed hypergeometric tests on the genes assigned to each enrichment group for every environment, separately, using the function “phyper” from the stats R package [44]. The resulting *P*-values were false-discovery rate adjusted, and significant results were plotted using ggplot2 to generate Figure 5C.

## RESULTS

**Table 1:** Sources of data used in this study.

Sources of data used in this study.
| Environment | Sample origin | Source | Virus enrichment approach | # of virome-metagenome sample pairs used | Sample design |
| --- | --- | --- | --- | --- | --- |
| Human gut | Fecal samples; Cork, Ireland | Shkoporov et al., 2019 [11] | 0.45 μm filtration, ultracentrifugation, & polyethylene glycol (PEG) precipitation | 10 | Individuals, timepoint |
| Freshwater | Oxic & anoxic water columns; Lake Mendota, Madison, WI, USA | Tran et al., 2023 [6] | 0.22 μm filtration & FeCl<sub>3</sub> precipitation | 14 | Water column depth, timepoint |
| Marine | Tara Oceans | Pesant et al., 2015; Sunagawa et al., 2015 [38,39] | 0.22 μm filtration & FeCl<sub>3</sub> precipitation | 21 | Water column depth, geographic location |
| Soil | Tomato field; Davis, CA, USA | Santos-Medellin et al., 2021 [27] | Amended 1% potassium citrate (AKC) resuspension, 0.22 μm filtration | 15 | Soil amendment, plot, timepoint |

### Viromes were successful in enriching for viral sequences

Sequencing depth within and between viromes versus metagenomes varied (Figure 2A). Freshwater and human gut viromes had a significantly higher sequencing depth than metagenomes, while marine metagenomes had a higher sequencing depth than viromes (Figure 2A). There was no difference in depth between viromes and metagenomes of soil samples (Figure 2A). Because of this observed variation in sequencing depth, results hereafter were normalized to sequencing depth unless otherwise specified. Reads from viromes of all environments mapped back to their assembled contigs (>10 kb) at a significantly higher rate than metagenomes (Figure 2B). Strikingly, soil viromes recruited upward of 25% of filtered reads while all soil metagenomes recruited less than <1% of filtered reads. Further inspection of soil metagenome assembly statistics revealed a median N50 <3,000, even when only calculating statistics for contigs >2,000 bp (Figure S1). The poor read recruitment of the soil metagenome assemblies is likely a result of the poor contiguity of the assemblies arising from high community complexity in soils [69,70].

Although the differences between viromes and metagenomes with respect to sequencing depth and read recruitment varied by environment, viromes from all environments had reads mapping to viral contigs at a greater rate than metagenomes (Figure 2C). All assemblies (metagenomes and viromes) except for the human gut had a greater proportion of viral to nonviral contigs (Figure 2D). Moreover, viromes from all environments except for the human gut had a higher total number of viral contigs than metagenomes (Figure S2A). Marine and soil viromes had a higher total number of vMAGs than metagenomes (Figure S2B). When considering only “high-quality” vMAGs that are estimated to represent complete or near-complete viral genomes [51], viromes from all environments had a greater yield than metagenomes (Figure S2C). Similarly, after dereplicating vMAGs to species-level clusters within samples, viromes had a higher viral species richness than metagenomes among marine and soil assemblies. However, there was no difference in viral species richness between methods among freshwater and human gut assemblies (Figure S2D).

### The abundance of lytic and lysogenic viruses in viromes *vs*. metagenomes varied

Among human gut assemblies, there was no significant difference between the number of lytic vMAGs from viromes compared to metagenomes, while freshwater, marine, and soil assemblies had a higher number of lytic vMAGs in viromes compared to metagenomes (Figure S3A). In contrast, there was no difference in the number of lysogenic vMAGs between viromes and metagenomes of freshwater and human gut assemblies, while marine and soil viromes contained significantly more lysogenic vMAGs than metagenomes (Fig S3B). Freshwater metagenomes contained significantly more vMAGs predicted to represent integrated prophage (Figure S3C). Integrated prophage vMAGs were found in viromes across all four environments (Figure S3C). Strikingly, marine and soil viromes contained significantly more integrated prophage vMAGs than metagenomes (Figure 3C). Closer inspection revealed that soil metagenomes did not contain any vMAGs predicted to represent integrated prophages at all. Given that the total number of vMAGs generated from marine and soil metagenomes was so low compared to their viromes (Figure S2B), these striking differences are explained by the low virus richness in these metagenomes overall. Last, while there was a small observable increase in the normalized number of integrated prophages in human gut metagenomes, these differences were not significant (Figure S3C).

### Viromes and metagenomes have unique and shared vMAGs

Dereplication and read mapping yielded 24,761 unique species-representative vMAGs in freshwater assemblies, 18,331 in marine assemblies, 9,039 in soil assemblies, and 2,271 in human gut assemblies, with a total of 54,402 unique vMAGs identified across all environments (Figure 3A). Of this total, 2,539 were found only in metagenome assemblies, 32,601 were found only in virome assemblies, and 19,262 were found in both (Figure 3B). Overall, virome assemblies from all four environments contained more unique vMAGs than metagenome assemblies (Figure 3C). Soil virome assemblies contained nearly all vMAGs detected in soil metagenomes, except for a single vMAG found unique to soil metagenomes (Figure 3C). Notably, more vMAGs were detected in both viromes and metagenomes of freshwater and human gut samples than were detected in either method, alone (Figure 3C).

We also examined the presence and absence of vMAGs in viromes and metagenomes separated by their predicted lytic state. More lytic vMAGs (Figure 3D), lysogenic vMAGs (Figure 3E), and integrated prophages (Figure 3F) were detected in viromes than metagenomes for all environments. However, freshwater assemblies had more lytic vMAGs detected in both methods than lytic vMAGs present in only one method (Figure 3D). Similarly, the human gut had more lysogenic vMAGs and integrated prophages present in both methods than those present in only one method (Figure 3E-F). However, the patterns of detection for integrated prophages may have been caused by virome reads originating from excised lysogenic/temperate virus genomes that had mapped to metagenome vMAGs integrated in host DNA.

### Virome assembly resulted in a more complete viral genome

Past arguments in favor of utilizing virome extractions to study viral communities have cited a tendency to assemble more complete viral genomes with greater depth than those assembled from metagenomes [21,27,60]. To test this, we identified the same species vMAG from a virome and from a metagenome. The virome-assembled viral genome was nearly 38 kb in length with 70 gene predictions (Figure 4, Table S2), and was predicted to be complete by CheckV [51] due to the presence of direct terminal repeats. The metagenome-assembled viral genome, however, was predicted by CheckV to be incomplete and was nearly 5 kb shorter than the virome assembly and contained only 57 gene predictions (Figure 4, Table S2).

The missing regions in the metagenome-assembled viral genome spanned both ends of the contig (Figure 4). These regions covered eleven genes with unknown functions that were present in the virome but not the metagenome assembly, as well as the first 527 bases of a phage portal protein (Figure 4, Table S2). Additionally, the virome-assembled viral genome contained a 130 bp region spanning two genes predicted to encode a hypothetical protein and a tail protein (Figure 4, Table S2). This 130 bp region was absent from the metagenome assembly, resulting in a single, fused gene prediction for a phage tail protein (Figure 4, Table S2). The only region we identified in the metagenome-assembled viral genome that was absent from the virome assembly was a single 3 bp sequence over the portal protein (Table S2). Finally, although this genome was incompletely assembled from the metagenome, metagenome reads mapped over the entire length of the virome-assembled genome (Figure 4, Table S3). Virome reads also mapped to both assemblies of the same genome with a depth up to two orders of magnitude greater than metagenome reads (Figure 4, Table S3).

### Viral genes are differentially abundant across viromes and metagenomes

We identified a total of 414,780 protein-coding viral genes after dereplication across all environments and extraction methods. Of these, 13,099 proteins came from human gut assemblies, 206,127 from freshwater assemblies, 116,900 from marine assemblies, and 78,654 from soil assemblies (Table 2, Table S4). Out of all dereplicated genes, a total of 72,082 unique genes were differentially abundant across extraction methods (Wald test *P* <0.05, FDR adjusted) (Table 2, Table S4). Only 55 of these genes were from the human gut, while 64,999 genes were from freshwater samples, 5,722 from marine samples, and 1,306 from soil samples (Table 2, Table S4). Using a minimum fold change cutoff of ±1.5, we found that 67,521 of the differentially abundant genes were enriched in either virome or metagenome samples (Table 2, Table S4, Figure 5A). The remaining 4,561 genes were differentially abundant but did not meet the minimum fold change of 1.5 (Table 2, Table S4, Figure 5A). We did not identify any genes that were enriched in either virome or metagenome samples from the human gut (Table 2, Figure 5A). However, 37,683 and 25,328 genes were enriched in viromes and metagenomes from freshwater samples, respectively (Table 2, Table S4, Figure 5A). Among marine samples, only 222 genes were enriched in viromes whereas 3,265 were enriched in metagenome samples (Table 2, Table S4, Figure 5A). Finally, 432 genes were enriched in soil viromes and 591 were enriched in soil metagenomes (Table 2, Table S4, Figure 5A).

**Table 2.** Number of genes throughout the differential abundance (DA) workflow.

| Environment | Number of genes before dereplication | Number of genes after dereplication (% of before) | Differentially abundant genes (% of dereplicated) | Virome-enriched genes (% of DA) | Metagenome-enriched genes (% of DA) |
| --- | --- | --- | --- | --- | --- |
| Human gut | $8.39 \times 10^4$ | $1.31 \times 10^4$ (16%) | 55 (0.004%) | 0 | 0 |
| Freshwater | $1.02 \times 10^6$ | $2.06 \times 10^5$ (20%) | $6.50 \times 10^4$ (32%) | $3.77 \times 10^4$ (58%) | $2.53 \times 10^4$ (39%) |
| Marine | $6.75 \times 10^5$ | $1.17 \times 10^5$ (17%) | $5.72 \times 10^3$ (4.9%) | 222 (3.9%) | $3.27 \times 10^3$ (57%) |
| Soil | $4.42 \times 10^5$ | $7.87 \times 10^4$ (18%) | $1.31 \times 10^3$ (1.7%) | 432 (33%) | 591 (45%) |
| **Total** | **$2.22 \times 10^6$** | **$4.15 \times 10^5$ (19%)** | **$7.21 \times 10^4$ (17%)** | **$3.83 \times 10^4$ (53%)** | **$2.92 \times 10^4$ (40%)** |

To predict potential functions for the differentially abundant genes enriched in either viromes or metagenomes, we used PHROG [68] functional categories predicted by Pharokka [63]. Out of the 67,521 unique genes enriched in viromes or metagenomes across all environments, Pharokka assigned PHROG functional categories to a total of 11,115 genes (16%), 6,247 in viromes and 4,868 in metagenomes (Table S4). Because predicted PHROG functional categories were largely present in both virome- and metagenome-enriched genes across the three environments (Figure 5B), we performed hypergeometric tests on enriched genes from each environment to determine whether any functional categories were over or underrepresented in viromes or metagenomes. We found nine PHROG categories that were significantly over- or underrepresented between viromes and metagenomes across freshwater, marine, and soil samples (hypergeometric test *P* <0.05, FDR adjusted) (Figure 5C, Table S5). Generally, genes encoding viral structural proteins such as head-tail connectors, packaging proteins, and tail proteins were underrepresented in metagenomes and overrepresented in viromes across freshwater and soil samples, while marine samples displayed the opposite pattern (Figure 5C, Table S5). Integration and excision coding genes were overrepresented in freshwater and marine metagenomes but underrepresented in freshwater viromes (Figure 5C, Table S5). Conversely, lysis genes were underrepresented in freshwater metagenomes and overrepresented in viromes, but were overrepresented in marine metagenomes.

## DISCUSSION

The sequencing of whole virus communities in recent years has resulted in an explosion of known viral diversity and viral community ecology studies [12,13,16,17,57,71]. Assembly of virus communities can be achieved either by sequencing extracted DNA from the total, mixed community of prokaryotes, eukaryotes, and viruses within a sample to generate metagenomes. Viral communities can also be assembled by enriching for virus-like particle DNA during extraction to generate viromes. Although viromes can generally offer a more focused view of viruses in a sample compared to metagenomes [33], the consequences of choosing one sampling method over the other have been relatively unexplored and limited to individual study ecosystems [5,6,27]. Here, we applied the same analytical methods to collections of paired virome and metagenome sequence reads to directly infer the unique and shared results gained from each sample method. We assembled, annotated, and analyzed 60 pairs of viromes and metagenomes across four different environments and found that the similarities and differences between each method varied across environments.

Viromes, by design, typically allow more viral species and genome coverage to be obtained compared to metagenomes [33]. In support of this, virome assemblies here generally contained more viral contigs, more binned vMAGs, higher species richness, and greater read recruitment to vMAGs. Interestingly, there were some exceptions among freshwater and human gut samples. We observed no difference in the number of vMAGs or in viral species richness between viromes and metagenomes of the human gut or freshwater. There was additionally no difference in the number of viral contigs from the human gut.

While there have been a handful of studies in the past that have examined viral community data resulting from viromes in comparison to metagenomes [6,11,27,72,73], even fewer have taken a closer look at specific genome-level differences that result across the two methods. While we only focused on one viral species in this context, we found that a virome assembly resulted in a more complete viral genome with greater sequencing depth than the genome assembled from a metagenome of the same sample. Notably, the metagenome sample contained reads that mapped over the entire length of the complete version of the genome. Although some viral genomes may be incompletely assembled in metagenomes, their full sequences may be assembled if the metagenome reads are mapped to a higher quality virome assembly or reference genome.

Freshwater and marine metagenome samples used here were recovered from >0.22 μm size fractions, while human gut and soil metagenomes were unfiltered by particle size. Considering this, any observed differences between viromes and metagenomes from freshwater and marine assemblies may have been driven by the approach used to generate the metagenomes. On the other hand, differences (or lack thereof) between viromes and metagenomes from soil and human gut assemblies may have been driven by the low abundance of viral DNA relative to nonviral DNA in bulk, unfiltered samples.

Nonetheless, both freshwater and marine metagenomes contained substantial numbers of viral contigs and vMAGs despite efforts to filter the viral fraction. Furthermore, there were striking differences between viromes and metagenomes from soil samples, as well as in human gut samples to a lesser extent, both of which did not have their viral fraction filtered from the metagenome fraction. Altogether, this highlights the importance of utilizing enrichment techniques that are tailored to the environment of interest and the research questions being asked.

Whether the purpose is to assign taxonomy [74], reveal mechanisms to avoid host defenses [75], identify auxiliary metabolic genes [76], or investigate mobile reservoirs for antimicrobial resistance genes [77,78], obtaining functional gene predictions is a critical step in analyses of viral communities. However, it can be quite challenging to assign functional predictions to viral genes annotated from metagenomic environmental data due to their large sequence diversity and the undercharacterization of viruses. Thus, annotating genes in complex viral communities often reveals a substantial amount of viral “dark matter” represented as genes with no known function that encode “hypothetical” proteins [23,79,80]. This challenge was indeed present here, as we could obtain functional predictions for only 16% of genes enriched in viromes or metagenomes. Nonetheless, we identified several functional categories across the three environments where genes were differentially abundant.

Our results show that one’s choice of extraction method does indeed influence the identification of gene families, but the significance and magnitude of differences vary between environments. We found an overrepresentation of integration and excision genes in freshwater and marine metagenomes with an underrepresentation in freshwater viromes. However, lysis genes were underrepresented in freshwater metagenomes and overrepresented in freshwater viromes. This is consistent with our observations that freshwater metagenomes contained a greater number of integrated prophage vMAGs than viromes. On the other hand, this contrasts with our observation that there was no difference in the proportion of lysogenic vMAGs between freshwater viromes and metagenomes, and that marine viromes contained more lysogenic and integrated vMAGs than metagenomes. Regardless of the exact mechanism(s), as a consequence, the choice between viromes and metagenomes can significantly influence one’s interpretation of viral communities based on gene annotations.

## CONCLUSIONS

In many contexts, viromes revealed more viral sequences and diversity than metagenomes. Hence, extracting viromes may be more advantageous than metagenomes when studying viral communities (Table 3). However, a noticeable number of viruses were detected only in metagenomes in all four environments tested here. Thus, we recommend that researchers investigating viral communities extract both viromes and mixed-community metagenomes in pairs from the same biological samples, when possible (Table 3). However, if one is restricted to using just one method, viromes present the better option for virus-focused studies in most environments.

**Table 3.** Recommendations for choosing extraction methods depending on research context.

| Context | Recommended method(s) | Rationale |
| --- | --- | --- |
| Viral community dynamics, overall virus diversity, assembly of uncultivable virus genomes | Virome | Viromes generally contained more viral species and greater viral sequence enrichment than metagenomes. |
| Bacterial/archaeal communities, no interest in viruses | Metagenomes | Viromes are unnecessary to the study of just the cellular members of communities. |
| Fast-growing, highly dynamic communities, and/or lytic viruses | Virome | Assuming viral lysis is prevalent due to the present biotic or abiotic conditions, viromes will enrich for lytic viruses. |
| Slow-growing, low-biomass communities, and/or integrated viruses | Metagenomes | Assuming lysogeny is prevalent due to the present biotic or abiotic conditions, detecting viruses integrated in the host genome require metagenomics. |
| Host-virus interactions | Paired viromes & metagenomes | Metagenomes are necessary to provide any host context. While metagenomes alone can yield some viral genomes, viromes are also recommended to maximize viral genome assembly. |
| Maximization of total virus diversity | Paired viromes & metagenomes | Both viromes and metagenomes resulted in the assembly of viral genomes not detected in the other method. Utilizing both methods can maximize the detection and assembly of as many viral genomes as possible. |

## DECLARATIONS

### Ethics approval and consent to participate

Not applicable.

### Consent for publication

Not applicable.

### Availability of data and materials

The datasets analyzed during the current study are available in the following repositories: Freshwater, originally presented by Tran et al. [6] and deposited to the JGI Genome Portal under Proposal ID <u>506328</u>; Marine, originally presented by Pesant et al. [38] and Sunagawa et al. [39] and deposited to the NCBI Sequence Read Archive under BioProject accessions <u>PRJEB1787</u> and <u>PRJEB4419</u>; Human gut, originally presented by Shkoporov et al. [11] and deposited to the NCBI Sequence Read Archive under BioProject accession <u>PRJNA545408</u>; Soil, originally presented by Santos-Medellin et al. [27] and deposited to the NCBI Sequence Read Archive under BioProject accession <u>PRJNA646773</u>. All scripts and intermediate files to reproduce the figures and tables presented here are available at <u>github.com/jamesck2/ViromeVsMetagenome</u>.

### Competing interests

The authors declare that they have no competing interests.

## Funding

This research was supported by National Institute of General Medical Sciences of the National Institutes of Health under award number R35GM143024, and by the National Science Foundation under grant numbers DBI2047598 and OCE2049478.

## Authors’ contributions

Conceptualization, J.C.K. and K.A.; Methodology, J.C.K. and K.A.; Software, J.C.K.; Validation, J.C.K.; Formal Analysis, J.C.K.; Investigation, J.C.K., K.M.K., M.V.L., P.Q.T., and K.A.; Resources, K.A.; Data curation, J.C.K., P.Q.T., and K.A.; Writing — Original Draft, J.C.K., K.A. ; Writing — Review & Editing, J.C.K., K.M.K., M.V.L., P.Q.T., and K.A.; Visualization, J.C.K.; Supervision, K.A.; Project Administration, K.A.; Funding Acquisition, K.A.

## Acknowledgments

We are thankful to all authors of the studies that originally generated and distributed the data analyzed here. We also gratefully acknowledge the insights provided by Cody Martin during this study.

## ABBREVIATIONS

VLP: Virus-like particle
ANI: Average nucleotide identity
PEG: Polyethylene glycol
vMAG: Viral metagenome-assembled genome
AKC: Amended 1% potassium citrate
vOTU: Viral operational taxonomic unit
DA: Differentially abundant

## ADDITIONAL FILES

**Additional file 1. Supplementary data and tables.** Includes Tables S1-S5 as referenced in the main manuscript text. File format: .xlsx.

**Additional file 2. Supplementary text and figures.** Includes supplementary methods, supplementary results, Figures S1-S6, and associated references. File format: .docx.

## Notes

### Competing Interest Statement

The authors have declared no competing interest.

### Summary of Updates

Main text, figures, and analyses have been updated. Specifically, we have updated analyses of marine viromes with new datasets and corrected some errors that were brought to our attention.

