## Supplementary Information for "Viromes vs. mixed community metagenomes: choice of method dictates interpretation of viral community ecology"

**SUPPLEMENTARY METHODS**

***Species accumulation curves***

Using multi-sample map files generated in the presence/absence analysis, CoverM v0.6.1 ([github.com/wwood/CoverM](https://github.com/wwood/CoverM)) was used to obtain trimmed-mean read coverage statistics (removing 5% of bases with highest coverage and 5% of bases with lowest coverage) for all species-representative vMAGs in each environment, as done by past investigations into virus presence/absence in community data [1,2]. These coverage tables were used with the R package vegan [3] with the function specaccum to generate to generate 1,000 random permutations of sample order using all tested sample pairs for each environment, separated by sampling method. Resulting cumulative species richness across all permutations, and their means, were plotted using ggplot2 [4] to generate Figure S6.

**SUPPLEMENTARY RESULTS**

***Obtaining paired viromes and metagenomes***

Our search for paired virome and metagenome datasets resulted in a collection of four different types of environments with a total of 120 samples (Table 1): a seasonally anoxic freshwater lake [5] sampled at discrete depths and timepoints, marine water columns from the global oceans sampled at discrete depths and geographic locations, the human gut microbiome in a cohort of ten healthy adults [6], and a dataset from a tomato agricultural field with spatially-structured soil samples across plots with varying soil amendments, which was previously used to directly compare viromes and mixed metagenomes in another study [7].

Viromes from all samples used size filtration to enrich for viruses, however, the exact approaches differed slightly between environments (Table 1). Viromes of freshwater, marine, and soil samples were generated by virion precipitation/resuspension and 0.2 μm filtration (Table 1) which is a widely used method to generate viromes [8]. Human gut samples underwent 0.45 μm filtration in addition to additional ultracentrifugation and polyethylene glycol (PEG) precipitation (Table 1) which has been established as optimal for extracting viromes from human fecal samples [9]. Furthermore, freshwater and marine metagenomes were sequenced from the >0.22 μm fraction of samples after filtering for virus-like particles (Figure 1). This results in DNA from prokaryotes, large viruses, and other microbes >0.22 μm. However, some virus-like particles <0.22 μm may remain in this fraction due to adhesion to cells or other surfaces [10] or because of clogged pores in the filter [11,12]. Soil and human gut metagenomes were sequenced from bulk samples that had not undergone filtration to separate the viral fraction (Figure 1). Although virome and metagenome extraction methods were not constant across the different environments, our main comparisons here are based on viromes versus metagenomes within the same environment, and thus was not determined to impact data interpretations and conclusions.

***Viral families shared and unique to viromes and metagenomes***

Having examined the degree of overlap in species-level vMAG clusters between viromes and metagenomes, we were interested in whether higher-level, defined viral taxa were uniquely found in viromes or metagenomes. Although only 17,776 out of all 54,402 unique representative vMAGs here could successfully be assigned to a described viral family (Figure S4), we used the previously generated presence/absence results in conjunction with taxonomy results generated by ViWrap (Figure 2A) to assess which known viral families were found in both viromes and metagenomes or unique to either environment.

Using the names of each viral family assigned to vMAGs from viromes and metagenomes, we found that both methods contained assemblies with many of the same viral families (Figure S5). Although 10,432 species-level vMAGs were unique to freshwater viromes compared to just 1,175 being shared with freshwater metagenomes (Figure 3C), 17 viral families were assigned to vMAGs from both freshwater viromes and metagenomes out of a total of 21 unique families assigned to vMAGs across both methods (Figure S5A). Steigviridae and Suoliviridae were unique to freshwater viromes, while Iridoviridae and Poxviridae were unique to freshwater metagenomes (Figure S4). All other environments had a complete overlap in the number of unique viral families assigned to vMAGs present in viromes and metagenomes, with 12 families assigned to marine vMAGs (Figure S5B), 11 families assigned to human gut vMAGs (Figure S5C), and 8 families assigned to soil vMAGs (Figure S5D).

Overall, of the vMAGs that could be assigned to a known viral family, identified viral families were generally present in both viromes and metagenomes. While there were two viral families unique to freshwater viromes and another two unique to freshwater metagenomes, the vMAGs assigned to these families were very rare relative to the other vMAGs in the environment (Figure S4). Since only a small fraction of vMAGs across all examined environments could be successfully assigned to a described virus family, there are likely many vMAGs within the same hypothetical “family” that were left out of the analysis shown in Figure S5. However, the near-complete overlap in vMAGs assigned to viral families known at the time of this analysis suggests that viromes and metagenomes may both contain a large proportion of vMAGs within the same virus family once virus taxonomy is expanded in the future.

***Sampling effort and viral species accumulation***

We asked whether reduced viral diversity observed in metagenomes could be alleviated by increasing metagenome sampling effort. Using 1,000 random permutations of sampling order, species accumulation curves showed varying cumulative richness across environments. Freshwater assemblies resulted in a higher cumulative richness in metagenomes than viromes, which remained consistent for all 14 sample pairs (Figure S6). In contrast, human gut, marine, and soil viromes consistently resulted in a higher cumulative richness than metagenomes for all sample pairs (Figure S6). Although, the human gut had cumulative richness curves suggestive of a significant number of undetected viral species in both methods (Figure S6). Lastly, increasing sampling effort up to 21 and 15 samples in marine and soil environments did little to bring the observed richness between viromes and metagenomes closer together, consistent with previous results of the same data [7]. The species accumulation curves from the four environments tested suggest that increasing the number of sample replicates has a large effect on observed viral species richness between viromes and metagenomes. However, the magnitude of the impact of sampling effort varies by environment.

**SUPPLEMENTARY FIGURES**

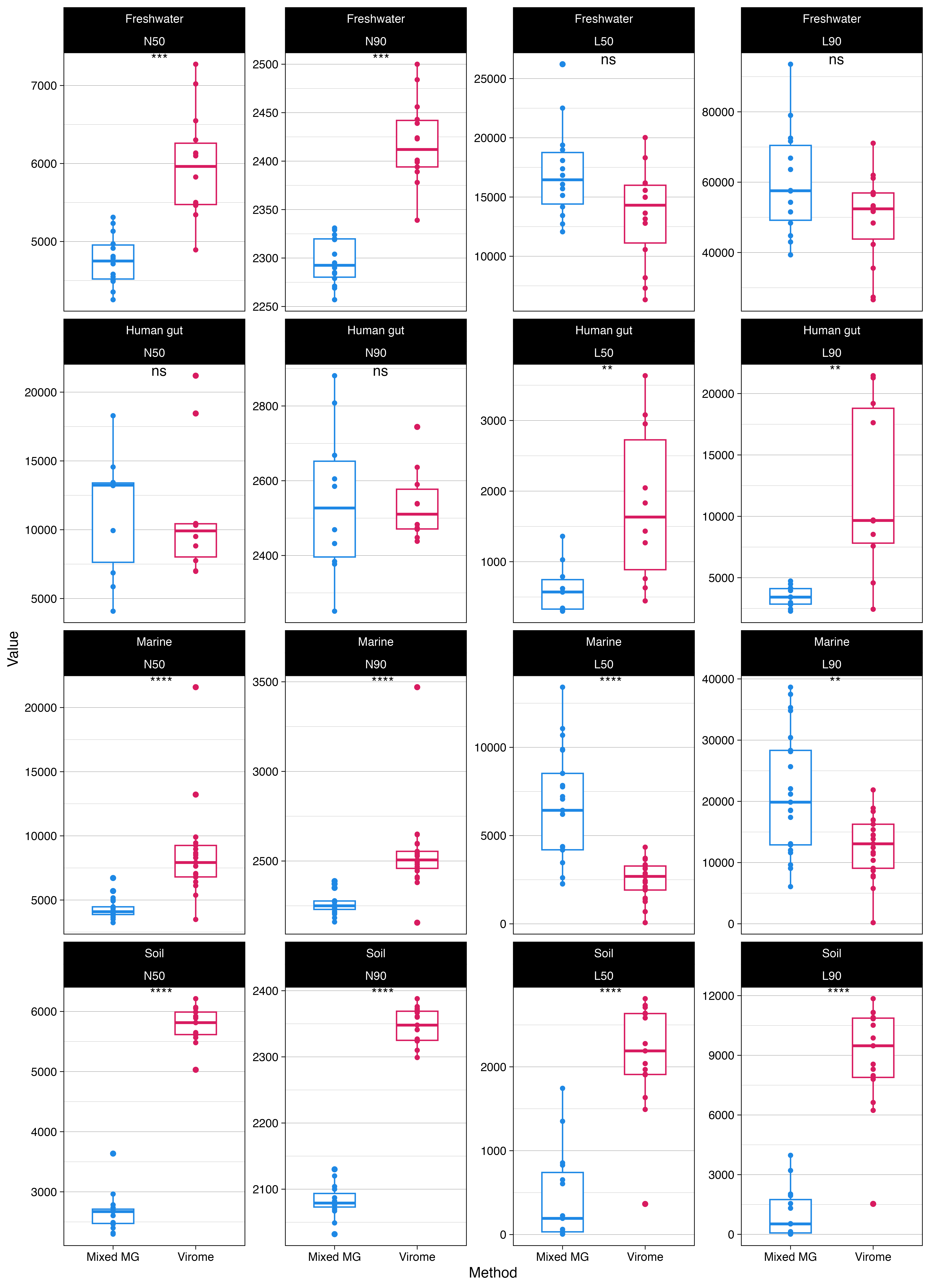

**Figure S1**. **Assembly statistics.** Points indicate an individual metagenome/virome assembly. All statistics were calculated only using contigs >2000 bp. Note that y-axes are not the same scale across panels. Significance was inferred by Wilcoxon rank sum test: ns *p* >0.05; * *p* ≤0.05; ** *p* ≤0.01; *** *p* ≤ 0.001; **** *p* ≤ 0.0001.

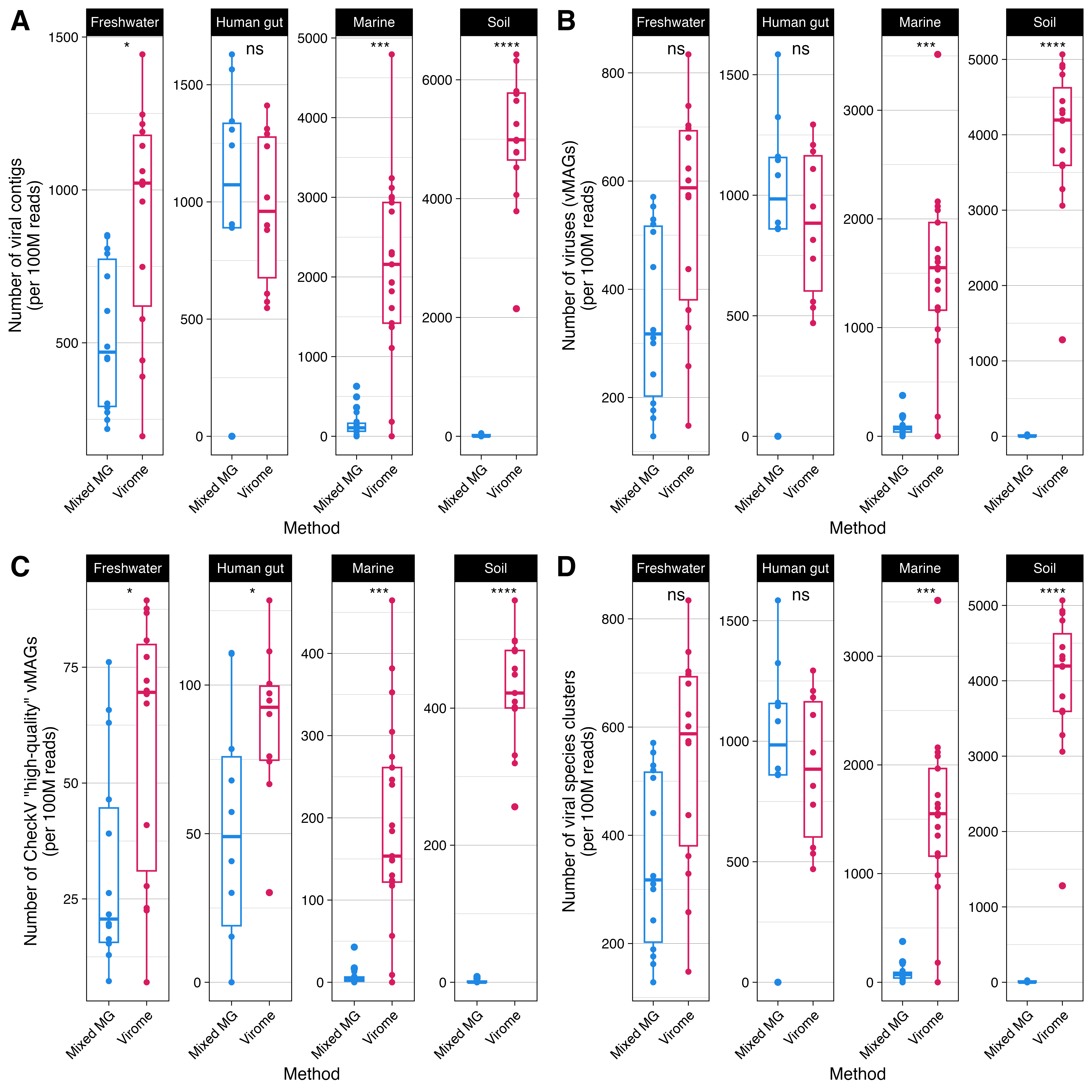

**Figure S2. Viromes generally outperformed metagenomes in all tested environments except for the human gut**. Results were normalized by dividing by the hundreds of millions of filtered reads per sample. Points indicate an individual metagenome/virome assembly. Note that y-axes are not the same scale across panels. Significance was inferred by Wilcoxon rank sum test: ns *p* >0.05; * *p* ≤0.05; ** *p* ≤0.01; *** *p* ≤0.001; **** *p* ≤0.0001. (A) Viromes resulted in significantly more viral contigs than metagenomes in all environments tested except for human gut samples. (B) Similarly, in marine and soil samples, viromes resulted in significantly more viral metagenome-assembled genomes (vMAGs) than metagenomes, whereas the number of vMAGs from freshwater and human gut samples was not different between viromes and metagenomes. (C) Among just high-quality vMAGs as determined by CheckV [13], viromes from all environments resulted in more vMAGs than metagenomes. (D) The number of viral species clusters at 95% average nucleotide identity was greater in marine and soil viromes, but not different between freshwater or human gut viromes and metagenomes.

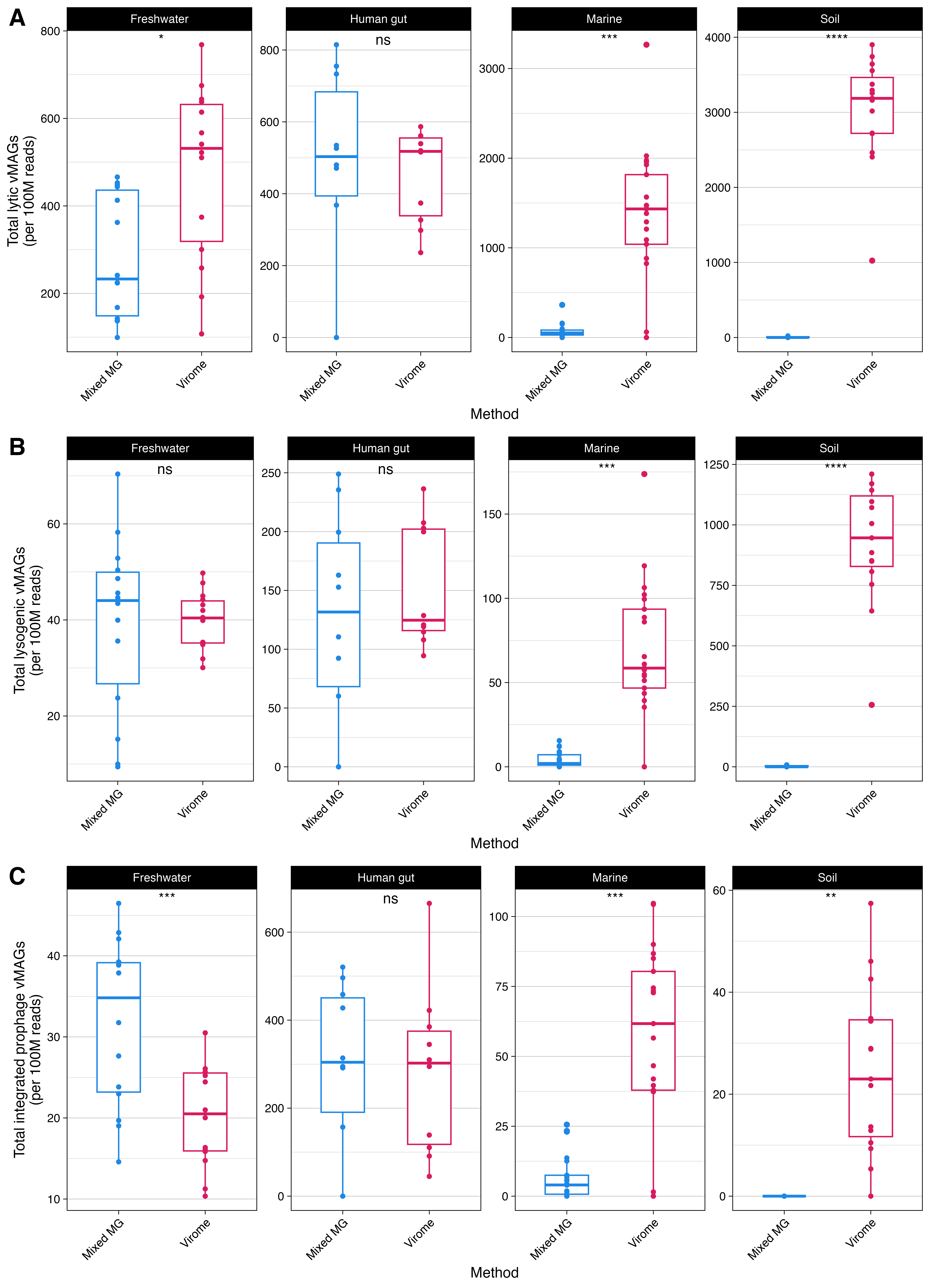

**Figure S3. The recovery of predicted lytic, lysogenic, and integrated viruses in metagenomes vs. viromes.** Points indicate an individual metagenome/virome assembly. Values represent the number of vMAGs in an assembly belonging to the facet category (i.e., lytic state ~ environment) normalized by hundreds of millions of filtered reads per sample. Note that y-axes are not the same scale across panels in (C). Significance was inferred by Wilcoxon rank sum test: ns *p* >0.05; * *p* ≤0.05; ** *p* ≤0.01; *** *p* ≤0.001.

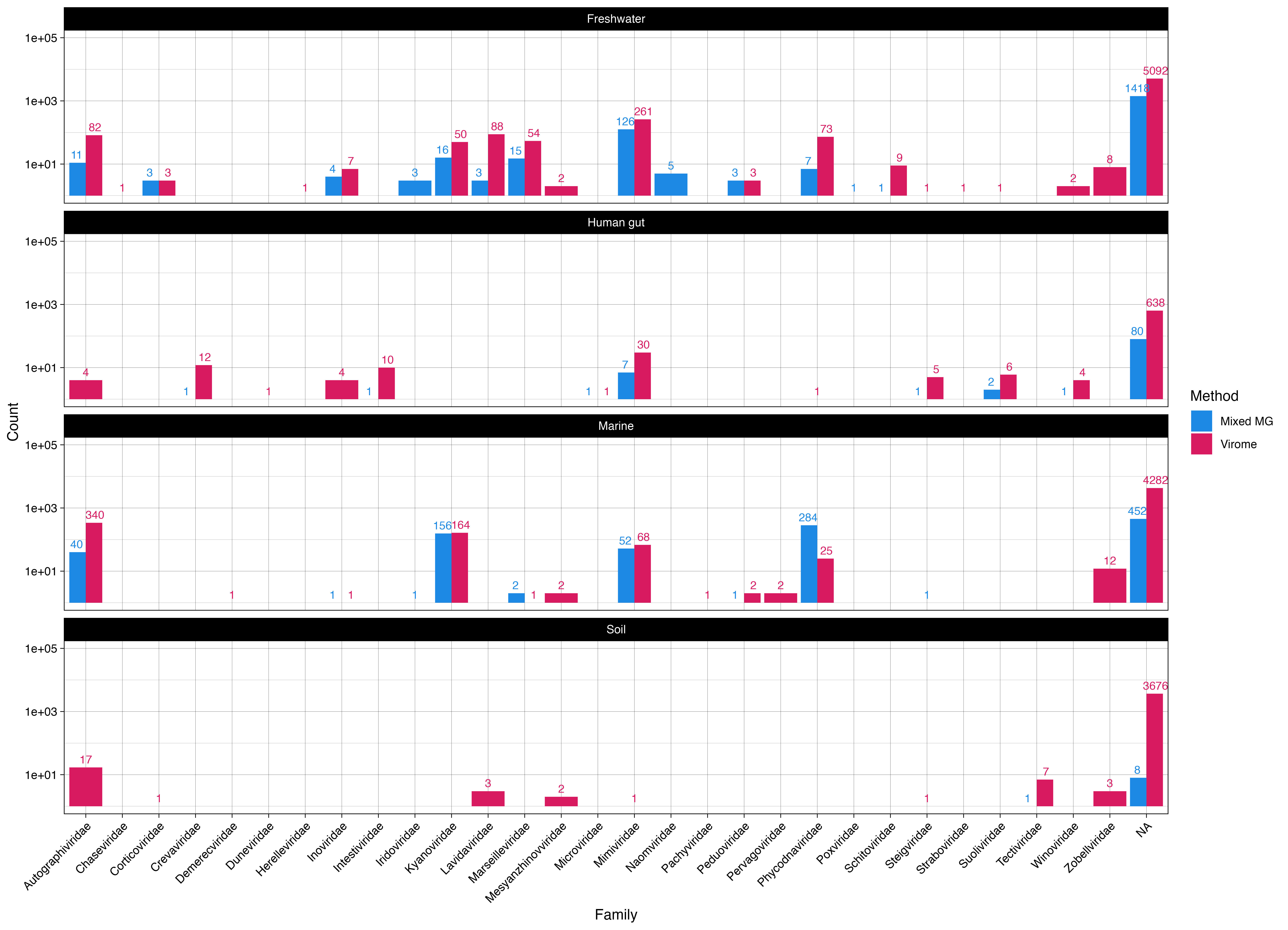

**Figure S4. Viral family assignments to species representative vMAGs.** Taxonomic assignments obtained from ViWrap pipeline [14]. Presence/absence of a vMAG in virome or metagenome assemblies determined by analysis summarized in Figure 1C.

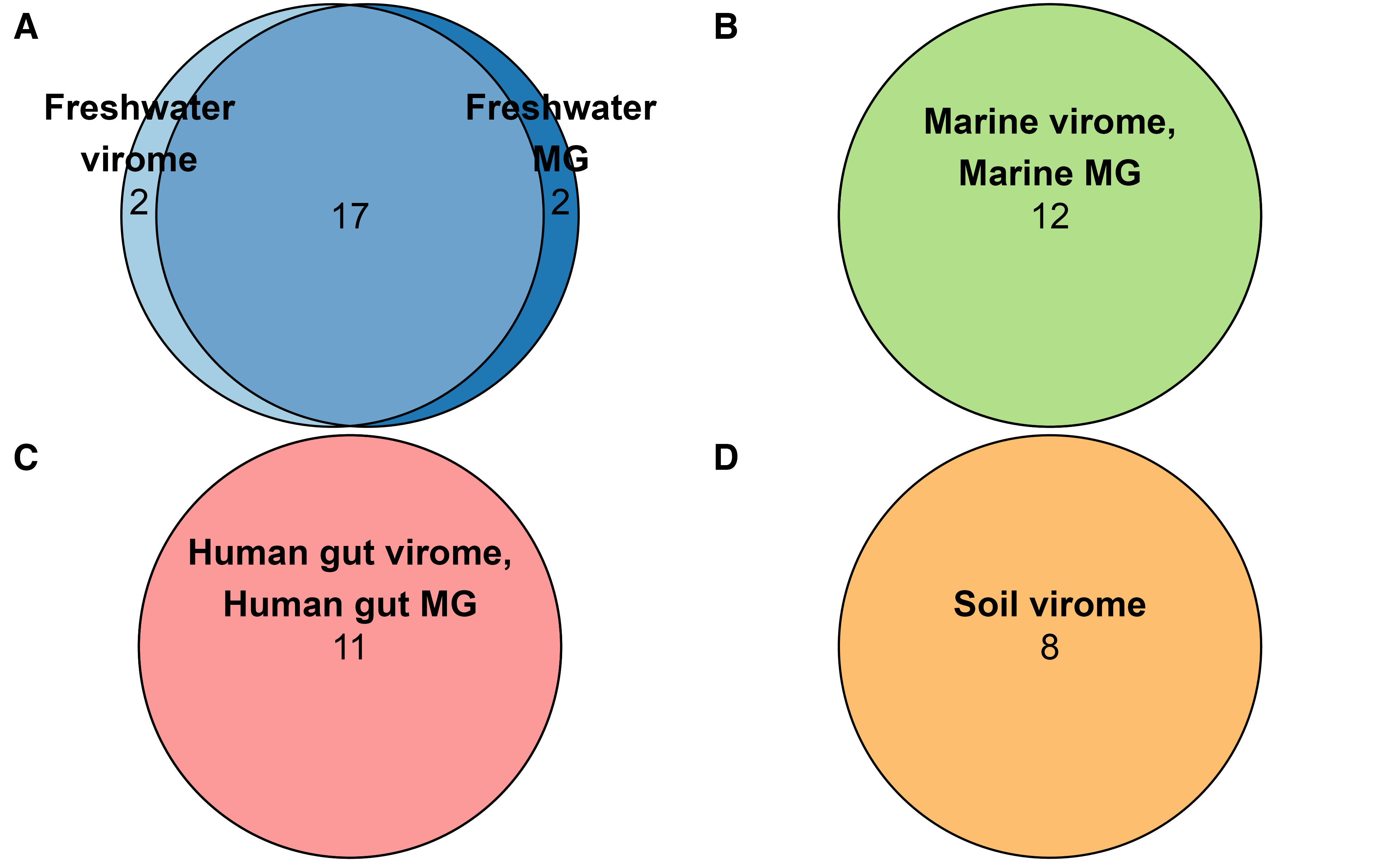

**Figure S5. Families of vMAGs found in viromes and metagenomes were largely found in both methods.** Euler diagrams generated using eulerr ([CRAN.R-project.org/package=eulerr](https://cran.r-project.org/package=eulerr)) [15,16] with names of unique, known viral families assigned to vMAGs detected in the labeled category; quantities within areas are given beneath labels. These Euler diagrams disregard overlap in families found in multiple environments and instead only considers the overlap in viral families found in viromes and metagenomes of (A) freshwater samples, (B) marine samples, (C) human gut samples, and (D) soil samples. Except for freshwater samples, there was a complete overlap in the unique viral families identified in viromes and metagenomes, with no known viral families identified in soil assemblies.

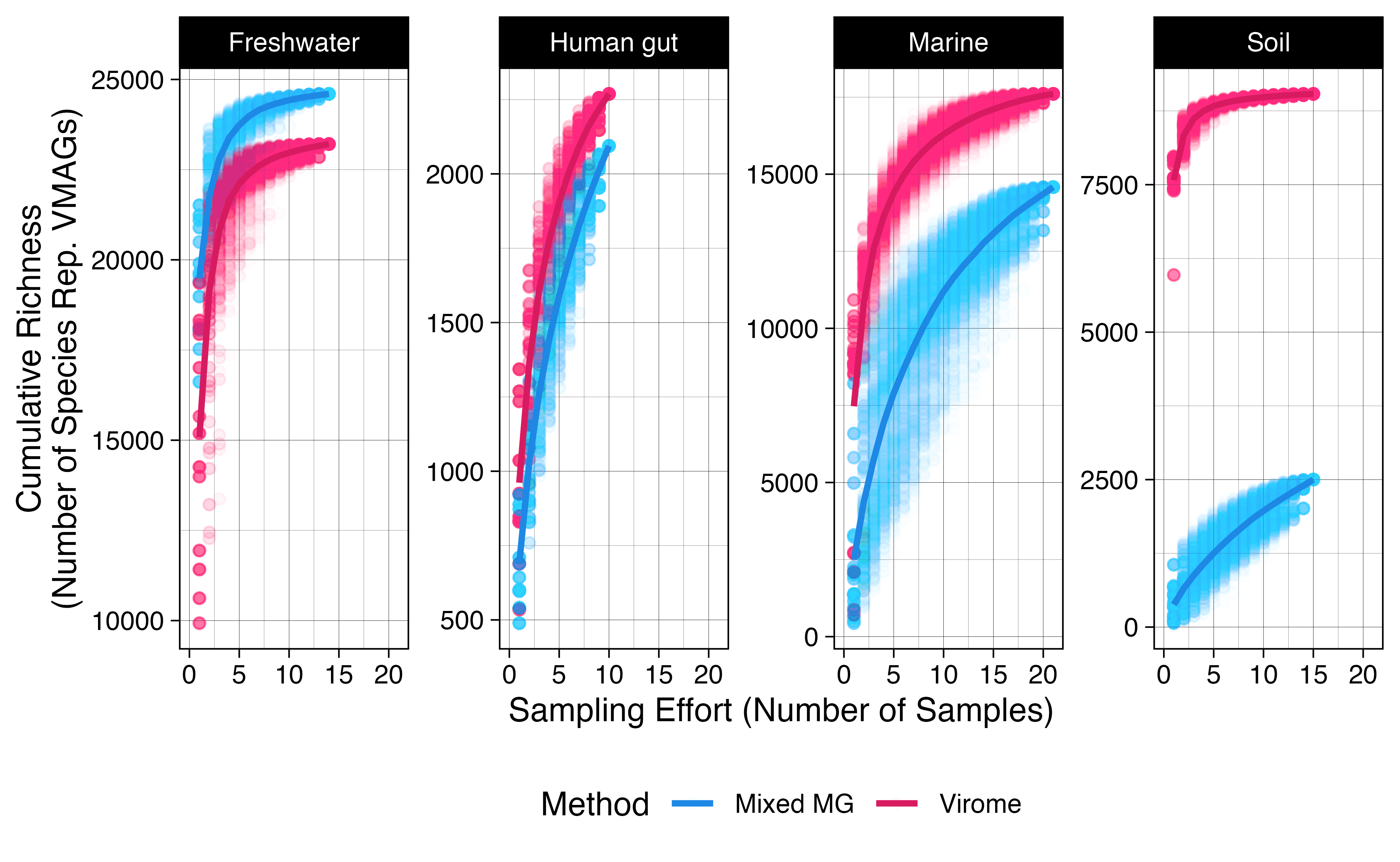
**Figure S6. The effects of sampling effort on cumulative viral species richness.** Points indicate the cumulative viral species richness across 1,000 random permutations of sampling order at each total number of samples along the x-axis. The average cumulative richness for each environment across all permutations for each sampling effort are represented by curves over points. The maximum sampling effort for each environment reflects the actual number for samples included in the analysis, with *n* = 14 for freshwater virome-metagenome pairs, *n* = 10 for human gut pairs, *n* = 21 for marine pairs, and *n* = 15 for soil pairs.
